# Spatio-temporal modeling of an insect vector distribution

**DOI:** 10.64898/2026.05.22.727173

**Authors:** Marie Grosdidier, Emily Walker

## Abstract

The pinewood nematode is a dangerous organism for the health of Pine forests and has already demonstrated its impact in Portugal and Spain. This organism was recently detected (November 2025) in France. It can spread over long distances (e.g., through wood transport) or short distances via dispersal of its airborne insect vector (about fifteen km), *Monochamus galloprovincialis*. The spatiotemporal distribution of the population of this insect vector is poorly understood in France. In this study, we propose to estimate the Monochamus population using official trapping data, which, however, are distributed heterogeneously in space and time. Spatiotemporal modeling is based on two complementary Bayesian approaches (an INLA-based generalized additive model, and a mecanistic-statistical approach), and integrates environmental covariates as well as laboratory-derived experimental data about the insect vector. The results provide estimates and predictions of the spatio-temporal distribution of *Monochamus galloprovincialis* in France. The species distribution varies according to environmental conditions and biological characteristics of the insect; the population dynamics thus can be estimated at all points in space and time. Prior to this study, no knowledge were available on the spatiotemporal distribution of the insect vector population; consequently, these results enable improvements to surveillance protocols. They will improve the pinewood nematode surveillance plan to better monitor free zones and buffer zones around the outbreak.

## 1 Introduction

The pinewood nematode (PWN), *Bursaphelenchus xylophilus* (Nickle 1970), originating from North America, attacks pines (*Pinus spp*.) and causes pine wilt disease (PWD). Later, it was first introduced in Asia (Japan) and then in Europe. As of April 2026, it is already present in Portugal and Spain, and was recently detected in southwest of France (on 3 November 2025 in the city of Seignosse (Folcher et al., 2025)). This nematode could be dispersed by two typical pathways: a long-distance dispersal pathway via contaminated wood materials (Castagnone et al., 2019; Haack, 2006; Robinet et al., 2011) and a short-distance dispersal pathway via insect vectors of the genus Monochamus (Coleoptera) (Robinet et al., 2011). *Monochamus galloprovincialis* is a Cerambycidae beetle endemic and widely distributed in Europe (https://inpn.mnhn.fr/espece/cd_nom/11779).

There are 127 species of Monochamus in the world but not all of them transmit pinewood nematode. In North America, Monochamus species capable of transmitting PWN to a tree, are *M. carolinensis, M. scutellatus*, and *M. mutator* (Akbulut and Stamps, 2012), and in Asia, mainly *M. alternatus*. In Europe, 5 species are present: *M. galloprovincialis, M. saltuarius, M. sartor, M. sutor* and *M. urussovi* (David, 2014), but only *M. galloprovincialis* is capable of vectorizing PWN (Sousa et al., 2001). *M. galloprovincialis* plays a key role in the invasion dynamics of pine wilt disease caused by the pinewood nematode and is the main vector of the nematode in Europe. European pines are sensitive to PWN and its vector. France has 2.7 million hectares of pines (16% of all forest areas and 5% of the entire French territory (source: http://www.pinsdefrance.com/tout-savoir-sur-les-pins/ressource-francaise-en-pin/)), and especially in the Landes forest which is a forest in southwestern France composed of monoculture maritime pines.

*M. galloprovincialis* completes its life cycle in one year on average at French latitudes. In warmer latitudes such as Portugal, it can complete two cycles per year (Frimino et al., 2017), and in colder latitudes, it can complete a single cycle every two years (Akbulut and Stamps, 2012). *M. galloprovincialis* lays eggs in dying pine trees (Aleppo pine, Maritime pine, and black pine in order of preference and more rarely in *Picea abies, Abies alba, Larix decidua* and occasionally *Cedrus spp*. (Fotini, 2008, Castagnone et al., 2019)). The wood-boring larvae develop (and acquire nematodes if the pines are contaminated) and emerge once the temperature conditions are favorable (late spring, early summer). Naves and Sousa, 2009 studied the temperature thresholds necessary for the development of *Monochamus galloprovincialis*, with 50% of the population emerging when the threshold of 822 degree-days is reached. Once out of the wood, the immature insect will find a healthy tree to make a first maturation meal, thus infecting healthy trees by depositing the pinewood nematode. Once mature, the reproduction phase occur, then egg-laying, thus completing the cycle. According to David et al., 2014 and David et al., 2017, *M. galloprovincialis* has an average flight capacity of 16 km over its entire lifespan, which is on average 126 days (min 50 to max 160 days of life) in a natural environment. Per day, the maximum flight capacity is 8.5 km for males and 6.8 km for females. The dispersal of *M. galloprovincialis* also varies depending on the environment. A heterogeneous landscape, with a mix of species, is a flight constraint for the insect, forcing it to modify its route (Nunes et al., 2021). Moreover, areas that have suffered forest fires are very attractive to Monochamus and promote the movement of individuals in these areas. These burned areas offer dying trees suitable for egg-laying and available healthy trees for the maturation meal in nearby unburned areas, which consequently increase Monochamus population (Lee and Kim, 2025, Jung et al., 2020).

The European Union (EU) has been monitoring the entry and establishment of certain pests for plants classified as priority quarantine organisms or OQPs (EU Regulation 2019/2072). Around twenty OQPs are currently selected by European experts who analyze the risks of entry and establishment for Europe (EFSA https://www.efsa.europa.eu/fr/topics/topic/plant-health and https://gd.eppo.int/standards/PM5/). PWN is an OQP. Each EU country is required to monitor appropriate surveillance systems to ensure that these OQPs have not entered or established on their territory or to detect the emergence of an outbreak as early as possible with a view to reacting quickly with management and control measures. Thus, the pinewood nematode and its vector are part of official surveillance plans mandated by the EU (see European and Organization, 2018 and Union européenne, 2016). According to the EU reglementation, pheromone traps are used to detect and monitor *Monochamus galloprovincialis*. Traps caught only a little part of the population. Torres-Vila et al., 2015 through their mark-release-recapture experiment show a trap catchability rate between 8 and 36% of the Monochamus population in Portugal. Robinet et al., 2019 also carried out a mark-release-recapture experiment in the Landes (southwest of France) and observed an average catchability rate of max 3% for mature beetles, similar to Nunes et al., 2021 who observed an average catchability rate of 2.2% [0 - 47%] in the same region.

Knowledge of Monochamus biology is often considered in models aimed at better understanding pine wilt disease and predicting the risk of disease entry, establishment, and spread (Togashi and Shigesada, 2006, Robinet et al., 2011, Robinet et al., 2019). For example, B. et al., 2018 uses in the parameters of his model the number of infected Monochamus emerging from an infected tree and available for dispersal. However, the population dynamics of Monochamus are not known. Species distribution models allow us to answer questions of population dynamics and to characterize the environment of these species.

Hierarchical models, and more precisely state-space models (Durbin and Koopman, 2012), allow us to establish a conceptual framework where: (i) the data layer concerns the probability distribution associated with the data knowing the model parameters and the latent ecological process; (ii) the latent variable is the hidden layer and corresponds to the ecological process studied; (iii) Hyperparameters control the latent process and the observations. To estimate the parameters and latent variables of this type of hierarchical model, it is often easier to implement Bayesian inference, which allows us to treat a series of simple problems linked together in a probabilistic way. This also allows us to take into account the uncertainty of the variables and parameters.

A spatiotemporal dimension can be added to these models, for example, using spatiotemporal random effects with the INLA (Integrated Nested Laplace Approximation) method, which is fast and easy to use and implemented in an R package. It uses latent Gaussian Markov random fields (GMRFs) to account for spatial dependence. Temporal dependence can also be added using various models, such as a first-order autoregressive model (Rue et al., 2017).

The objective of this work is to estimate the population dynamics of *Monochamus galloprovincialis* in France through two complementary Bayesian modeling approaches, (i) INLA as described above, and (ii) a mechanistic-statistical model, both based on trapping data of *M. galloprovincialis* carried out as part of official surveillance for the pinewood nematode, and taking into account the biology and environment of the beetle.

## 2 Material & Method

### 2.1 Official monitoring data

France has been monitoring the insect vector, *M. galloprovincialis*, since 2012. Traps (Crosstrap®) diffusing pheromones (Galloprotect®) are set during the Monochamus flight period (between approximately April and November) in areas with pine trees and close to at-risk sites. The trap has an attractive radius of 100m (Jactel et al., 2019) and is set on average 10 days before being checked. These traps only catch mature adults (pheromones) and kill the insects with the presence of an insecticide. The agents in charge of monitoring come to collect the insects in the traps and send them to laboratories to analyze the presence or absence of the pinewood nematode (Mariette et al., 2023). This data is then pooled by the Ministry of Agriculture and Food Sovereignty (DGAl) and the ESV Platform.

The traps are not located on a regular grid and are not surveyed regularly. They can be relocated throughout the year, and are rarely reset on the same date and in the same location from one year to the next. In total, the traps were set at 2,753 sites in France and 9,517 trapping surveys were carried out between 2012 and 2024. Thus, the data collected are irregular in space and time; the sampling effort (i.e. time of trap setting) is different for each data collected.

### 2.2 Variables : description and selection

Variables related to hosts (number of trees, surface area, health status, quantity of dead wood), disturbances (fires, forest losses), land use and weather conditions (temperature, precipitation, emergence phenology, relative humidity) were calculated from different data sources and at different spatial scales around the traps. Three main categories of variables were selected by Random Forest, highlighting the importance of each variable in explaining the data. Experts from the pinewood nematode (PWN) Working Group of the ESV Platform also validated this selection, which overlaps with knowledge from the literature (see supplementary information).

First, the presence of host trees and their spatial configuration is taken into account. The surface area of pine trees attractive to *M. galloprovincialis* and sensitive to the nematode (Aleppo pine, Maritime pine, Scots pine, Corsican and Black pine) was calculated from the Forest Database V2 (IGN) over a 100m radius around the trap. Host surface data vary from 0 to 0.5 km^2^ for a 100m radius around the trap. To take into account the variability of the environment around the trap, a mixing rate is estimated from the surface area of host pines compared to other trees recorded in an area of 7500m around the trap (mean between female and male dispersal distances from David et al., 2014). This rate varies between 0 and 1, 0 corresponding to a pure forest of host pines and 1 to a forest without host pines, and with all intermediates.

Then, the weather emerges as a sine qua non condition for the analysis of these captures. The weather data (temperatures and liquid precipitation, named precipitations in the following) come from the SAFRAN reanalysis from Météo France (https://meteo.data.gouv.fr/datasets/donnees-changement-climatique-sim-quotidienne/). For each grid cell and each year, the cumulative daily average temperature above 12.2°C is calculated. The days when this cumulative temperature reaches 450, 608, 822, 1029, and 1146 allow the emergence dates of approximately 1, 10, 50, 90, and 99% of the Monochamus population to be determined, respectively (Naves and Sousa, 2009). For each trapping period, the maximum temperature and the sum of precipitations were calculated. Some maximum temperatures in France averaged over the trapping period remain negative. This corresponds to areas where the insect cannot develop. These periods are removed from the dataset and analysis.

Finally, the attractiveness of *M. galloprovincialis* for burned areas is evident in the pre-selection of covariates. Forest fire data are downloaded from the EFFIS/WILDFIRE Database (https://forest-fire.emergency.copernicus.eu/) by choosing the MODIS Burnt area product from 2011 to 2024. The distance from each trap to the nearest area composed of conifers that had burned the year before the trap was set (n-1) was calculated.

### 2.3 Validation and prediction

Official monitoring data from 2012 to 2021 were used to calibrate the two models (estimation step). Data from 2022 to 2024 were used to validate (validation step) the outputs.

To estimate the Monochamus population across France from 2022 to 2024 and correct for trap exposure in the prediction step, traps were artificial set every 10 days throughout the year to comply with the protocol initially defined by the state services, and an 8 x 8 km grid (calibrated to Météo France’s SAFRAN grid) was chosen. Moreover, two covariates have been normalized as follows for compatible scales with observed data. To estimate the pine area according to an attractivness of 100m around a hypothetical trap, a cross product was calculated on the pine area of the total quadrat. Similarly, to estimate mixture species rate according a radius of 7500m around a trap, a cross product was calculated with mixture species rate of the total quadrat.

### 2.4 INLA model

The first approach uses the R package INLA (Lindgren et al., 2011; Rue et al., 2017) and allows to take into account the spatial and temporal dependencies expressed in the data while estimating the results with a very short computation time.

After several tests (Poisson/negative binomial distribution, other covariates, different meshes) and a selection using the best Deviance Information Criterion (DIC), the model that emerges as the best one is based on a negative binomial distribution adapted to an overdispersed distribution of counts including many zeros but also many high values. The response variable *Y*_*zyt*_ corresponds to the number of Monochamus captured by a trap at site *z* =1, …, *Z* in year *y*= 1, …, *Y* on survey date *t*=1, …, *T*.

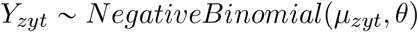

with

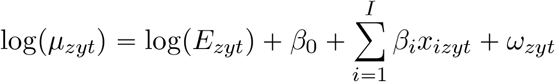

where *µ*_*zyt*_ and *θ* are respectively the mean and the dispersion parameters of the negative binomial distribution; *E*_*zyt*_ is a known exposure term (trap exposure time) integrated as an offset for observation zyt, *β*_0_ is the intercept, and *β*_*i*_(*i* = 1, …, *M*) are the fixed effects related to the covariates *x*_*i*_(*i* = 1, …, *M*).

The term *ω*_*zyt*_ refers to the residual spatiotemporal process capturing the spatiotemporal variability not explained by the environmental covariates *x*_*i*_, modeled using the INLA-SPDE approach. The variables *x*_*i*_ are MixSpecies rate 7.5km2, MaritimePine 100m, AleppoPine 100m, Corsican-BlackPine 100m, ScotsPine 100m, TMax, Sprecip, Dist min fire (Table 1). The construction of the mesh used to discretize space for the spatiotemporal effect is detailed in supplementary information. The prior model is a spatiotemporal Gaussian process, structured with first-order autoregressive (AR(1)) dynamics and spatially correlated according to a Matérn correlation function.

**Table 1:** Description of covariates.

| Variable | Description |
| --- | --- |
| MixSpecies_rate_7.5km2 | Mixing rate within a radius of 7.5 km around the trap. 1 corresponding to a forest without pines, 0.5 to a mixed forest between pines and non-pines and 0 to a pure pine forest |
| MaritimePine_100m | Area of maritime pines within a radius of 100 m around the trap |
| CorsicanBlackPine_100m | Area of Corsican or Black pines within a radius of 100 m around the trap |
| ScotsPine_100m | Area of Scots pines within a radius of 100 m around the trap |
| AleppoPine_100m | Area of Aleppo pines within a radius of 100 m around the trap |
| TMax | Maximum temperature over the trapping period |
| SPrecip | Sum of precipitation over the trapping period |
| Dist_min_fire | Minimum distance to the nearest coniferous forest fire in year n-1 |
| Allpine | Sum of MaritimePine_100m, CorsicanBlackPine_100m, ScotsPine_100m, AleppoPine_100m |

Spatial variability is taken into account by a dedicated parameter in the model. Thus, the areas of each host pine species could be integrated into the regression, unlike the mechanistic-statistical model where they were aggregated into a host pine variable (see below).

### 2.5 Mecanistic-statistical model

This second approach uses the R package Nimble (de Valpine et al., 2025) with a model defined by the combination of a latent layer representing population dynamics modeled deterministically within an annual period (inspired by the work of Soulsby and Thomas, 2012), and an observation layer modeled stochastically. This hierarchical model is composed of the following three embedded steps.

### Observation process

This hierarchical model is based on an inhomogeneous Poisson process. The number of trapped Monochamus Y, for a trapping period *t*_*p*_ for each site z and year y, is modeled as follows:

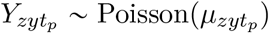

The observation process is taken into account in the intensity *µ*_*zyt*_*p* of the Poisson point process. Indeed, the intensity will depend on the sampling effort (trapping time *t*_*p*_ = *t*_*survey*_ − *t*_*setting*_), the abundance dynamics *λ*_*zyt*_*p*, a trap catchability parameter *ν*_1_, as well as the climate, and more specifically precipitation (with temperature already taken into account in the emergence function defined below) (Table 2).

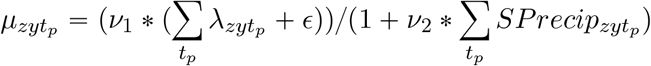

**Table 2:** Emergence function parameters.

| % emergence | Time interval | Slope | $E'(t)$ value |
| --- | --- | --- | --- |
| 0-1 | before $t_{0zy}$ | 0 | 0 |
| 1-10 | $t_{0zy} - t_{1zy}$ | $\alpha_{1zy}$ | $N_{1zy} * \alpha_{1zy}$ |
| 10-90 | $t_{1zy} - t_{2zy}$ | $\alpha_{2zy}$ | $N_{1zy} * \alpha_{2zy}$ |
| 90-99 | $t_{2zy} - t_{3zy}$ | $\alpha_{3zy}$ | $N_{1zy} * \alpha_{3zy}$ |
| 99-100 | after $t_{3zy}$ | 0 | 0 |

**Table 3:**
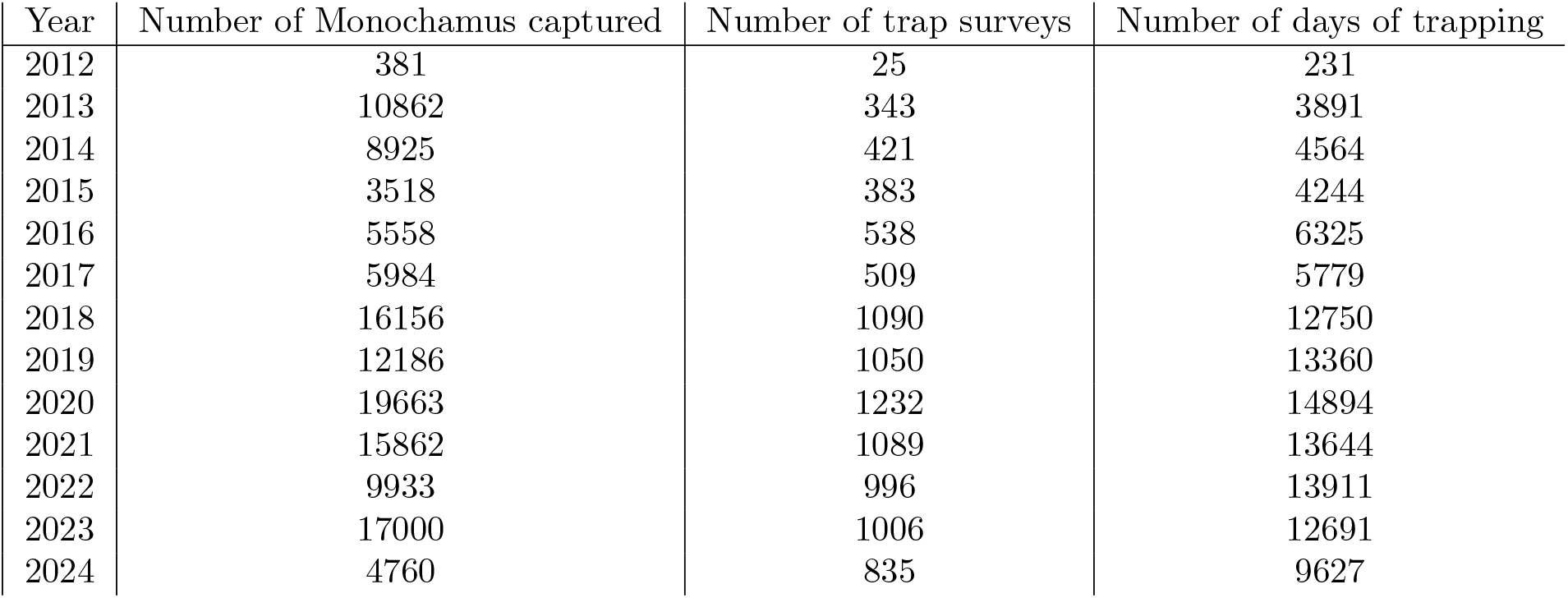
Number of Monochamus captured and number of trap setting per year.

| Year | Number of Monochamus captured | Number of trap surveys | Number of days of trapping |
| --- | --- | --- | --- |
| 2012 | 381 | 25 | 231 |
| 2013 | 10862 | 343 | 3891 |
| 2014 | 8925 | 421 | 4564 |
| 2015 | 3518 | 383 | 4244 |
| 2016 | 5558 | 538 | 6325 |
| 2017 | 5984 | 509 | 5779 |
| 2018 | 16156 | 1090 | 12750 |
| 2019 | 12186 | 1050 | 13360 |
| 2020 | 19663 | 1232 | 14894 |
| 2021 | 15862 | 1089 | 13644 |
| 2022 | 9933 | 996 | 13911 |
| 2023 | 17000 | 1006 | 12691 |
| 2024 | 4760 | 835 | 9627 |

For identifiability reasons, *ν*_1_ is set to 0.02 (Nunes et al., 2021). Setting *ϵ* = 0.000001 avoids division by 0. The parameter *ν*_2_ associated to precipitations will be estimated by inference.

### Mecanistic model

Given *ρ* the mortality rate and *D* the lifespan, the intra-annual population dynamics *λ*_*zyt*_, i.e. the Monochamus number for each year *y*, site *z* and day of year *t*, is written as:

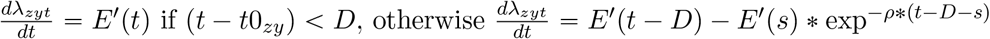

This dynamics is split into two terms:

- an emergence deterministic function *E*^1^p*t*q which takes different values depending on the time t as detailed in (Table 2).
with *t*0_*zy*_, *t*1_*zy*_, *t*2_*zy*_, *t*3_*zy*_ emergence dates for 1, 10, 90, 99% respectively; N1 the population available;
- and a mortality function:

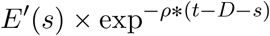

with *ρ* the mortality rate set at 4.75 (calculated from the survival curves of David et al., 2017) and *D* the lifespan set at 126 days (David et al., 2014).

### Regression on environmental variables

For each site z and year y, *N*_1_ is calculated as follows:

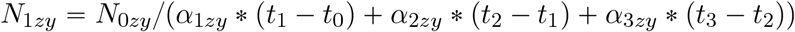

where *N*_0*zy*_ is the maximum daily available population over the year y for site z, expressed as:

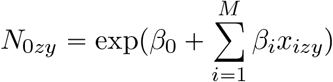

with *β*_0_ intercept and *β*_*i*_ fixed effects related to covariates *x*_*i*_.

Variables *x*_*i*_ are MixSpecies rate 7.5km2, Allpine, Dist min fire (Table 1).

The mechanistic-statistical model was parameterized for a single annual generation (Akbulut and Stamps, 2012). Since this approach does not account for spatial variability, host pine areas are considered globally (sums of all species combined) to avoid bias. Since spatial variability is not taken into account in the model, host pine areas were aggregated into a single host pine variable.

### Inference

The model is estimated with a Bayesian MCMC algorithm, running three independent chains for 20,000 iterations, with the 1,000 first iterations removed for burn-in, and a thinning factor of 10 to reduce autocorrelation in posterior samples. Weakly informative priors were used for the model parameters to be estimated (*β, ν*_1_) (see supplementary information). The calculations were run on a high-performance computer cluster.

## 3 Results

### 3.1 Descriptive statistics of data

9,517 surveys were realized in total between the days of year (DOY) 109 and 350. No Monochamus were captured in 26.17% of the surveys (2,491 surveys). The average number of Monochamus captured per day can range from 1 to 32. The number of trapping days varies between 1 and 158 days, with a median of 11 days and a mean of 12 days. The months with the highest average Monochamus captures per survey were June and July, followed by August and September (Figure 1). More Monochamus were captured in relatively warmer latitudes (southern France, latitudes below 46°5’) (Figure 2).

**Figure 1:**
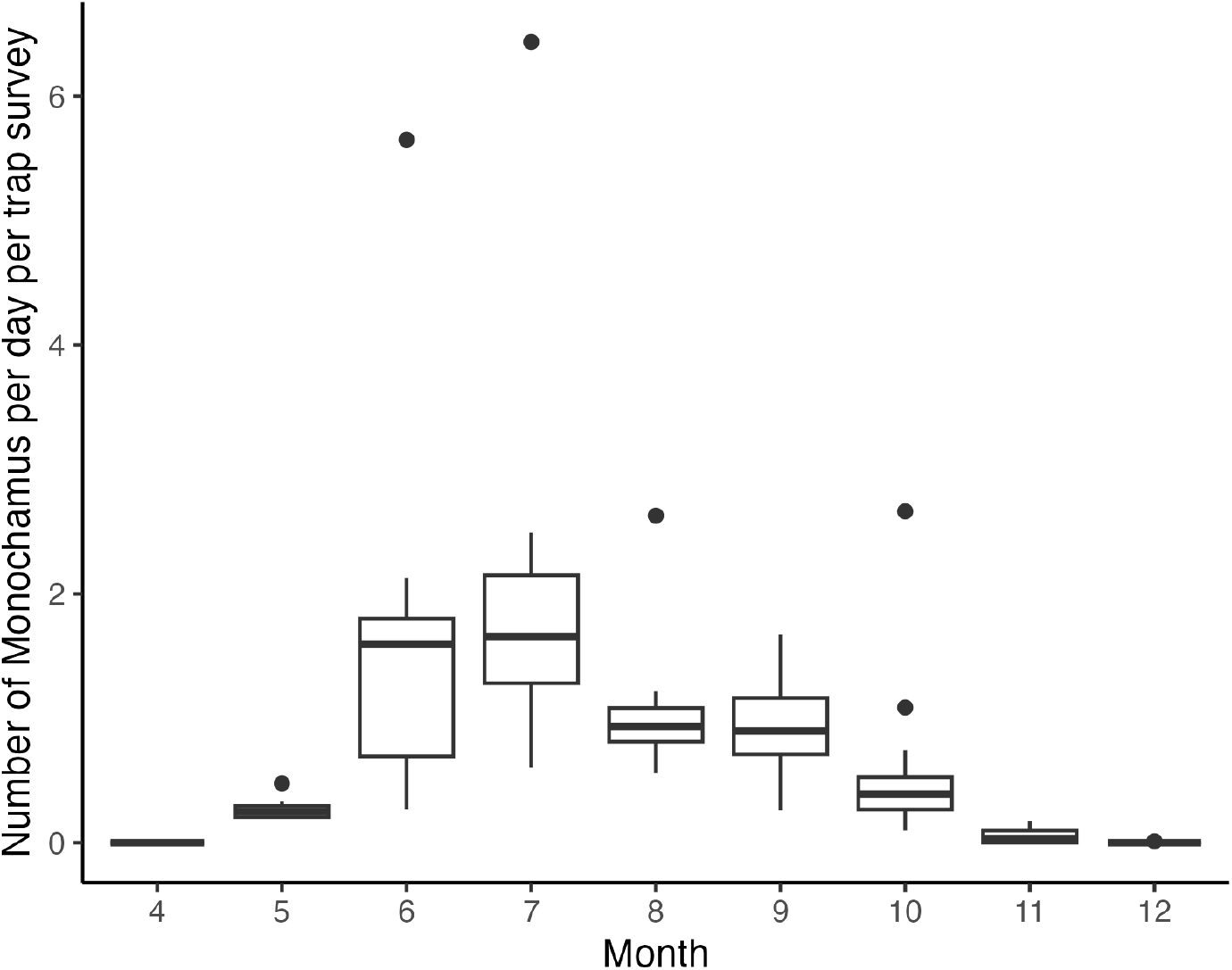
Intra-annual variability of observed data : number of Monochamus captured per day per trap setting according month of the year

**Figure 2:**
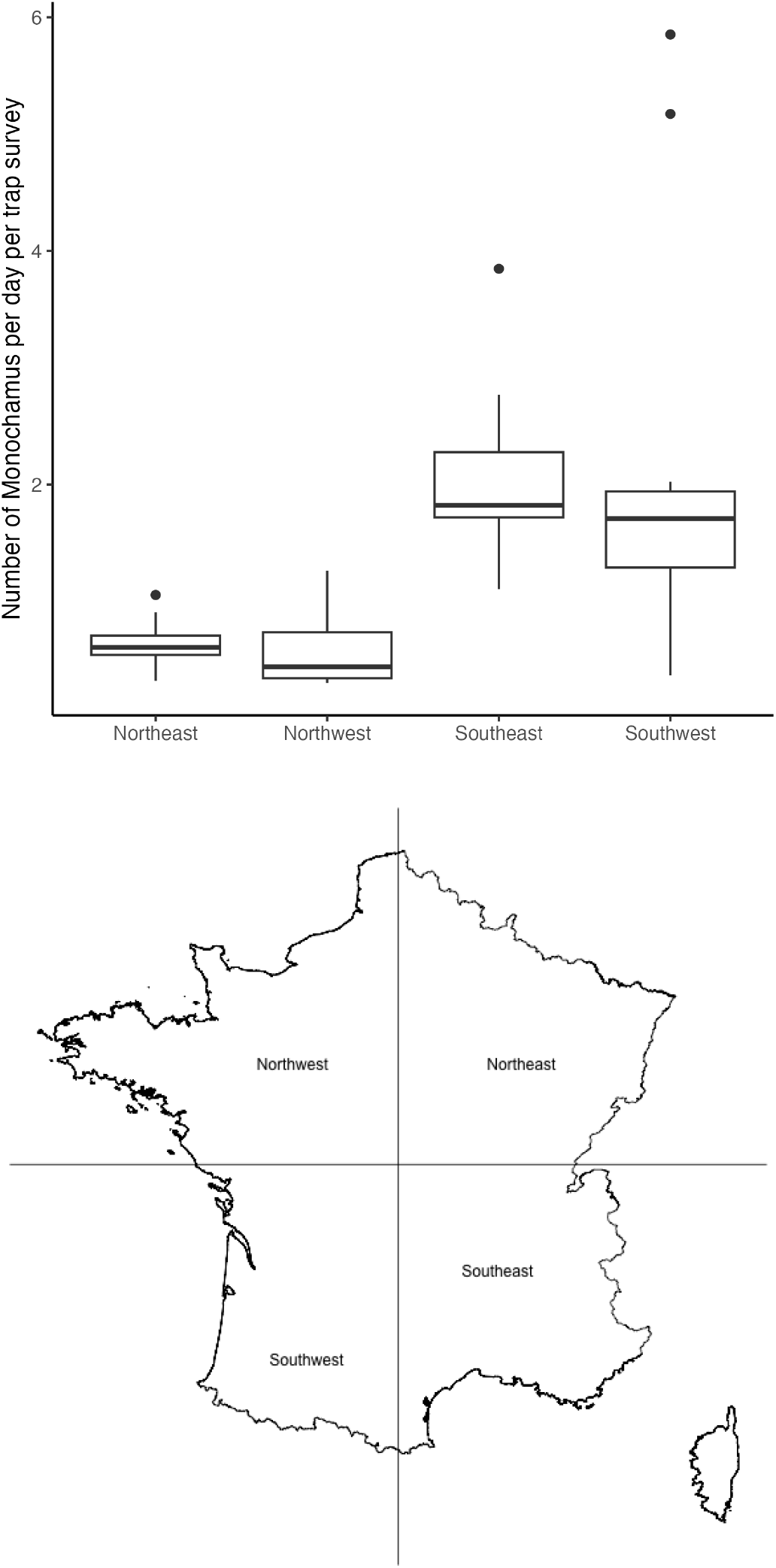
Spatial variability of observed data : number of Monochamus captured per day per trap setting arbitrary divided into 4 zones (with longitudinal division at 2.41666 and latitudinal division at 46.53333 (WGS84)

**Figure 3:**
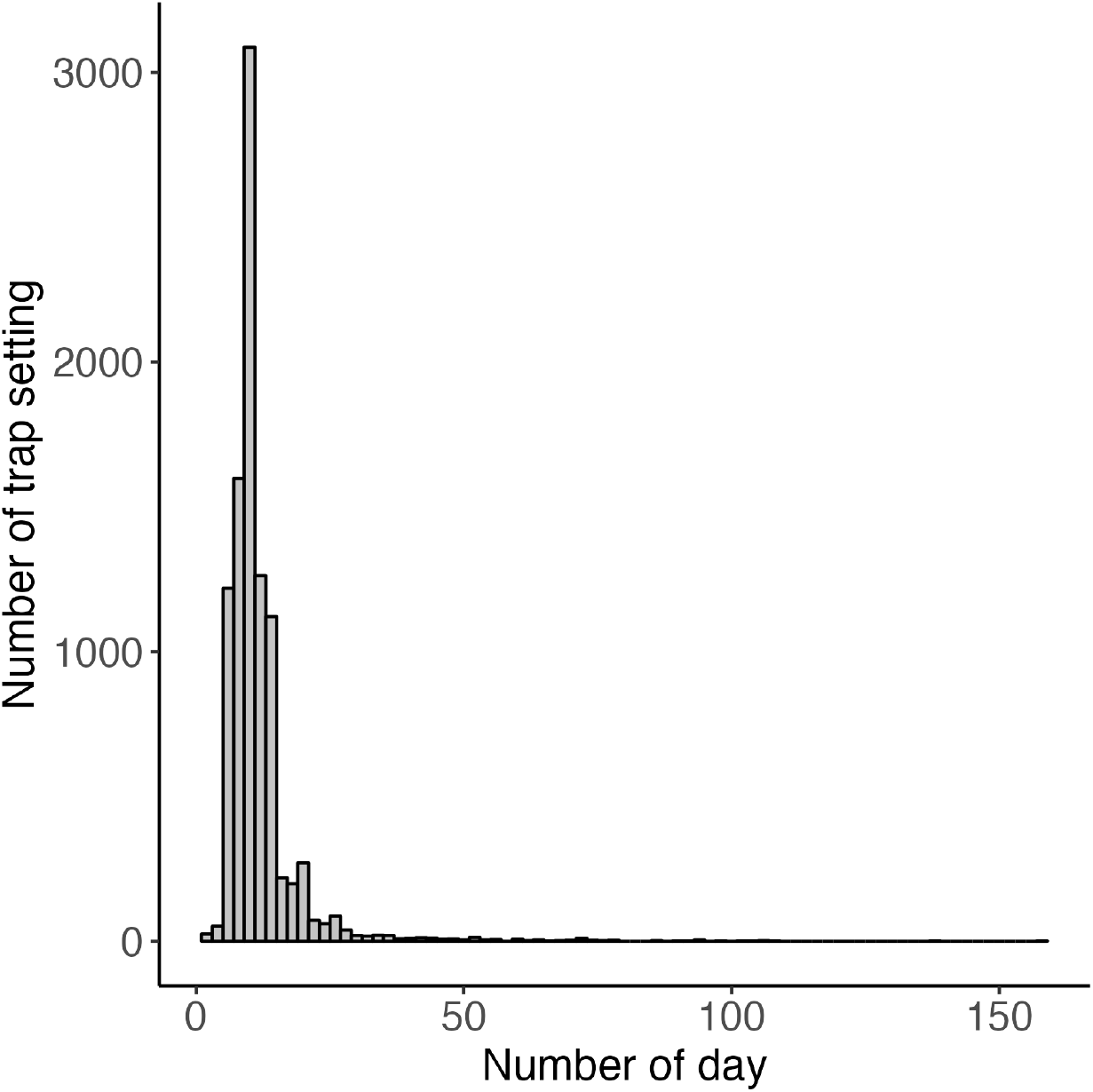
Temporal variability of trapping duration (in days)

Over the whole study period (2012-2024), the first and the last days with captured Monochamus in northeast of France are DOY 146 and 309 respectively, in northwest of France DOY 152 and 317, respectively, in southeast of France DOY 139 and 297, respectively, and in southwest of France DOY 126 et 339, respectively.

### 3.2 INLA model

From the covariates and data from 2012-2021, the model was used to estimate the parameters related to each covariate (step (1)). Then in a second validation step (2), the model was used to estimate a number of Monochamus for the site-years zy 2022-2024 (estimates compared to the data observed for the same site-years). In a third step (3), the model was used to predict the number of Monochamus at the national territory scale per quadrat for the years 2022 to 2024 according to a time step of 10 days.

#### 3.2.1 Estimation

The best model is the negative binomial model with a DIC of 39,223 (compared to 53,118 for the Poisson distribution). The overdispersion parameter is 1.18 (Table 5), indicating rather strong overdispersion. This indicates that strong overdispersion remains in the residuals after covariates and spatiotemporal random effects have been taken into account. Therefore, the variability of data with a large number of observed zeros (when there are no captures) and heavy-tailed numbers of Monochamus per trap can only be partially explained by the latent predictor of the model.

The parameters estimated by the model (Table 4) are all significant (the value 0 is not included in the quantiles). The signs are consistent with results from the existing literature. The mixing rate has a negative impact on the number of Monochamus captured, i.e. the more one is in a pure pine forest, the more Monochamus is captured. Each pine species has a positive impact on the number of Monochamus captured. The model with the normalized data allows us to prioritize the importance of pine species in the capture of Monochamus: Maritime pine > Black Corsican pine > Aleppo pine > Scots pine (see supplementary information). High precipitation during the trapping period negatively impacts Monochamus captures, and conversely, high maximum temperatures during the trapping period positively impact Monochamus captures. Proximity to a forest fire from the previous year positively impacts the number of Monochamus that can be captured.

**Table 4:** Model parameter results (unnormalized data)

| Parameter | mean | sd | 0.025quant | 0.975quant |
| --- | --- | --- | --- | --- |
| Intercept | -13.03 | 0.53 | -14.07 | -12.00 |
| Mix species rate | -2.20 | 0.39 | -2.95 | -1.44 |
| Aleppo pine | 43.41 | 7.07 | 29.55 | 57.26 |
| Corsican Black pine | 25.56 | 4.66 | 16.42 | 34.70 |
| Scots pine | 14.52 | 4.74 | 5.22 | 23.80 |
| Maritime pine | 19.53 | 3.22 | 13.21 | 25.85 |
| Precipitation | -0.08 | 0.01 | -0.11 | -0.06 |
| Temperature | 4.36 | 0.12 | 4.13 | 4.59 |
| Distance to fire | -0.10 | 0.03 | -0.17 | -0.04 |

Regarding the spatio-temporal aspect (Table 5), the estimated temporal autocorrelation coefficient (GroupRho), very close to 1, highlights strong interannual correlation between the data. Furthermore, a spatial correlation of approximately 60 km, unexplained by the model covariates, is observed, which corresponds to a regional scale. The size for negative-binomial observations is very small assuming no zero inflation.

**Table 5:**
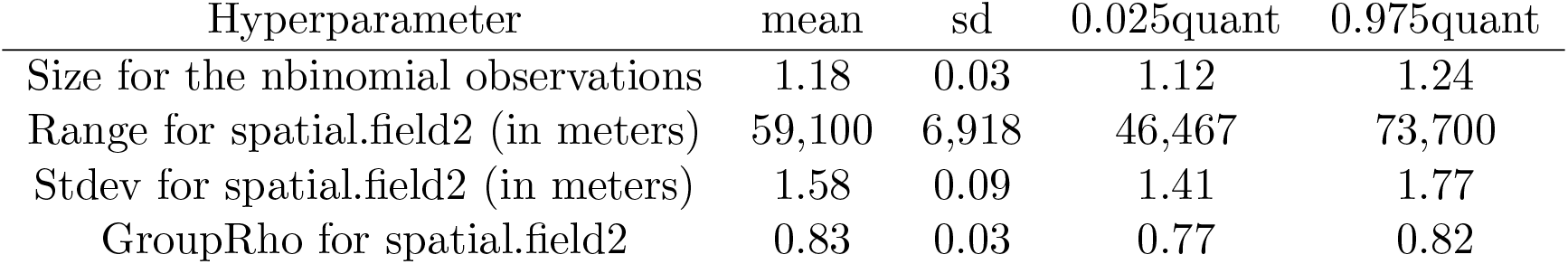
Model parameter results (unnormalized data)

The spatial field (Figure 4) shows an area along the English Channel where Monochamus is almost never caught. In the rest of France, the results show patches of Monochamus abundance that vary in size depending on the year.

**Figure 4:**
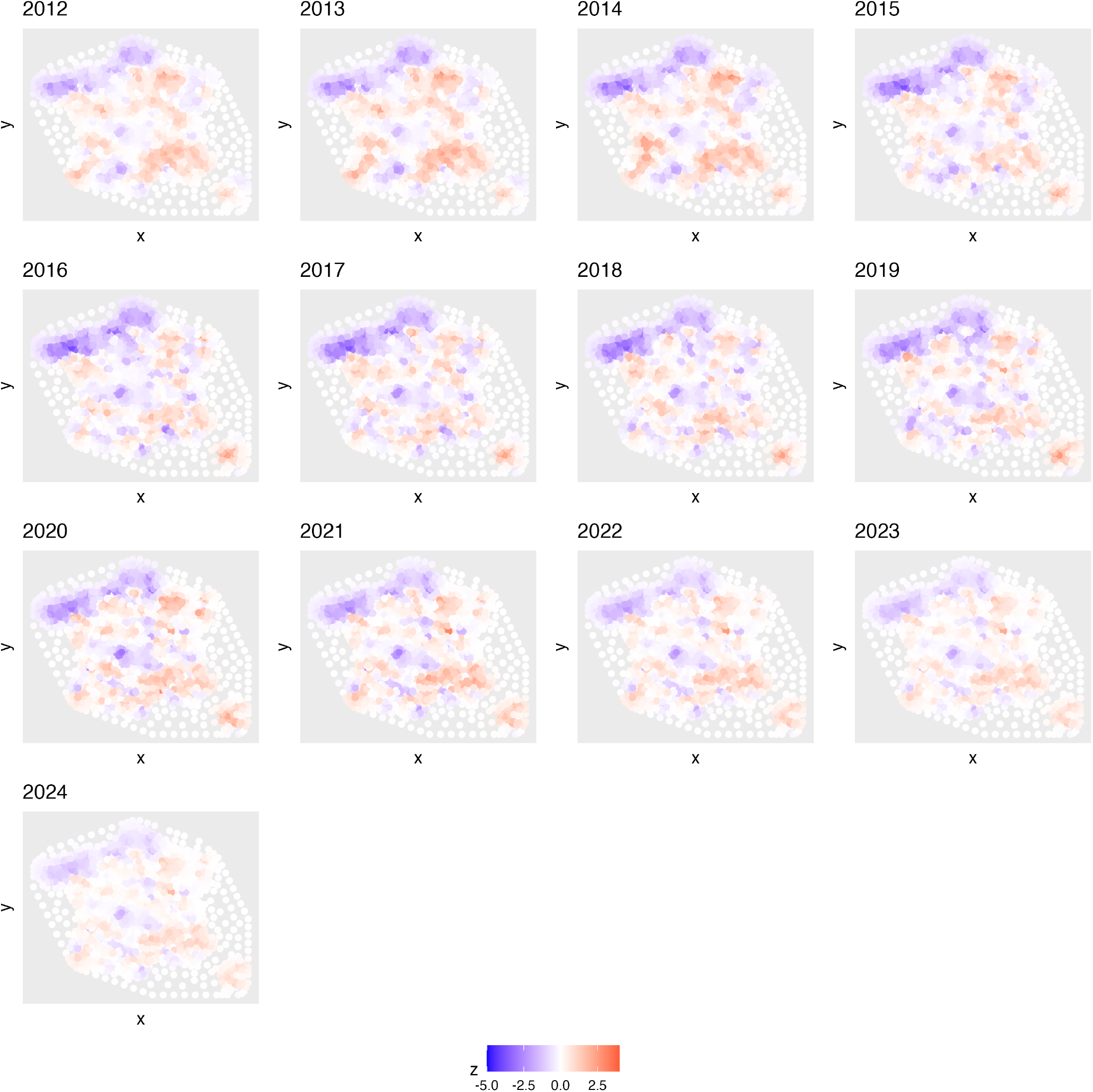
Annual spatial effects that is not explained by covariates across years

#### 3.2.2 Validation

We see that it overestimates the number of Monochamus captured but still remains acceptable with an R^2^ of 0.74 for data from 2012 to 2021 (estimation step) and 0.40 for data from 2022 to 2024 (validation step) (Figure 5).

**Figure 5:**
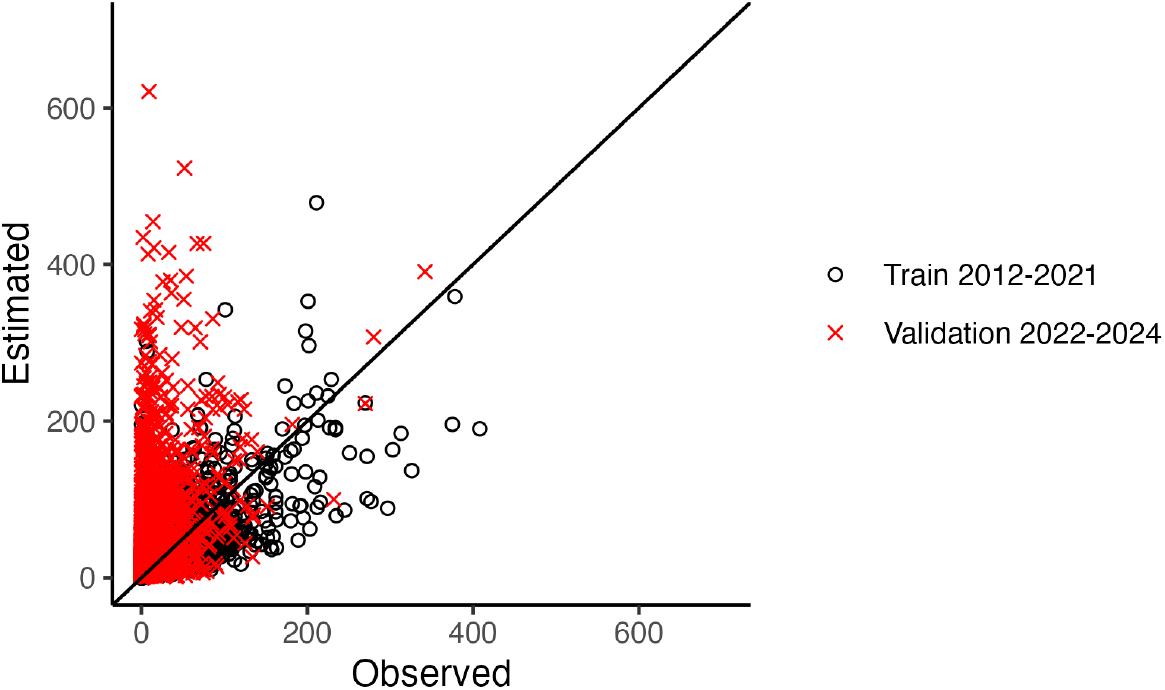
Number of Monochamus estimated and observed in 2022 to 2024. Each point may have different trapping time, but it remains consistent between observed and estimated values.

The model seems to take into account temporal variability well, both intra and inter annual; the trends follow each other (Figure 6). In the validation data, the first day with detected monochamus is around 142-168 and the last day with detected monochamus is around 265-335 depending on the geographical location. This corresponds very well with the observed data (first day around 143-168 and last day around 265-327). But, it is less good at estimating the quantity of Monochamus that can be caught. Model clearly overestimated number of Monochamus catchable in validation step 2022-2024 (Figure 6).

**Figure 6:**
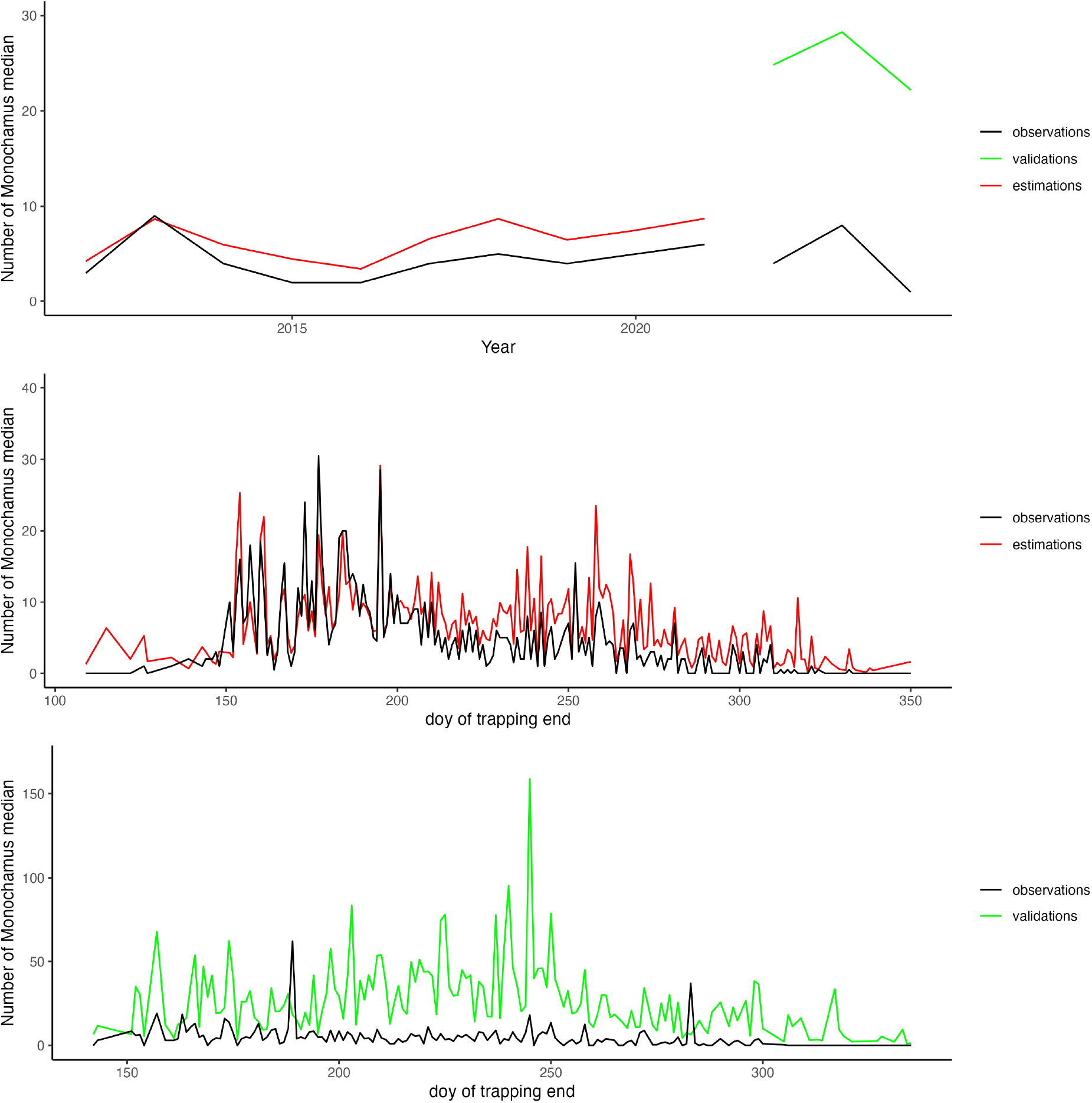
Temporal variability according observed data and INLA model recalculated values (top: inter-annual variability, middle: intra-annual variability for train data (2012-2021), down: intraannual variability for validated data (2022-2024)

#### 3.2.3 Predictions

The number of Monochamus that can be caught in each quadrat for the years 2022 to 2024 was estimated for 10-day trapping periods. Figure 7 shows a median representation of the 3 years (2022, 2023, 2024) of prediction of the sum of Monochamus that can be captured per month. According to the predictions, Landes forest and south-east of France would be areas where some Monochamus (less than 15) could be capture along all the year. And these are also areas where the amount of catchable Monochamus could be very important. On contrary, the areas along the English Channel show almost all year round either a catchable Monochamus population close to zero or with a number of fewer than 10 catchable individuals per month. Figure 8 shows the median catchable Monochamus population predicted temporally for 10 days of trapping for the 3 years according north-east, north-west, south-east or south-west location in France. We observe that the Monochamus population begins to emerge around day 125, with a few small captures possible around the first 100 days. Slight inter-annual variability is observed, but very little variation is observed at the geographic level in terms of dates. The geographic area’s influence is primarily in the quantity of Monochamus that can be captured (the south allows for the capture of more Monochamus). Finally, there appear to be no more Monochamus that can be captured globally after day 325, or even day 300 for the year 2023. If we focus on quadrats precisely distributed in different areas of France and the one containing the focus of the nematode detected in Seignosse in 2025 (red dot), as well as Figure 9, we see a strong spatial variability in the quantity of Monochamus that can be captured between each site, but also temporal variability with emergence dates and end of population that can be slightly offset.

**Figure 7:**
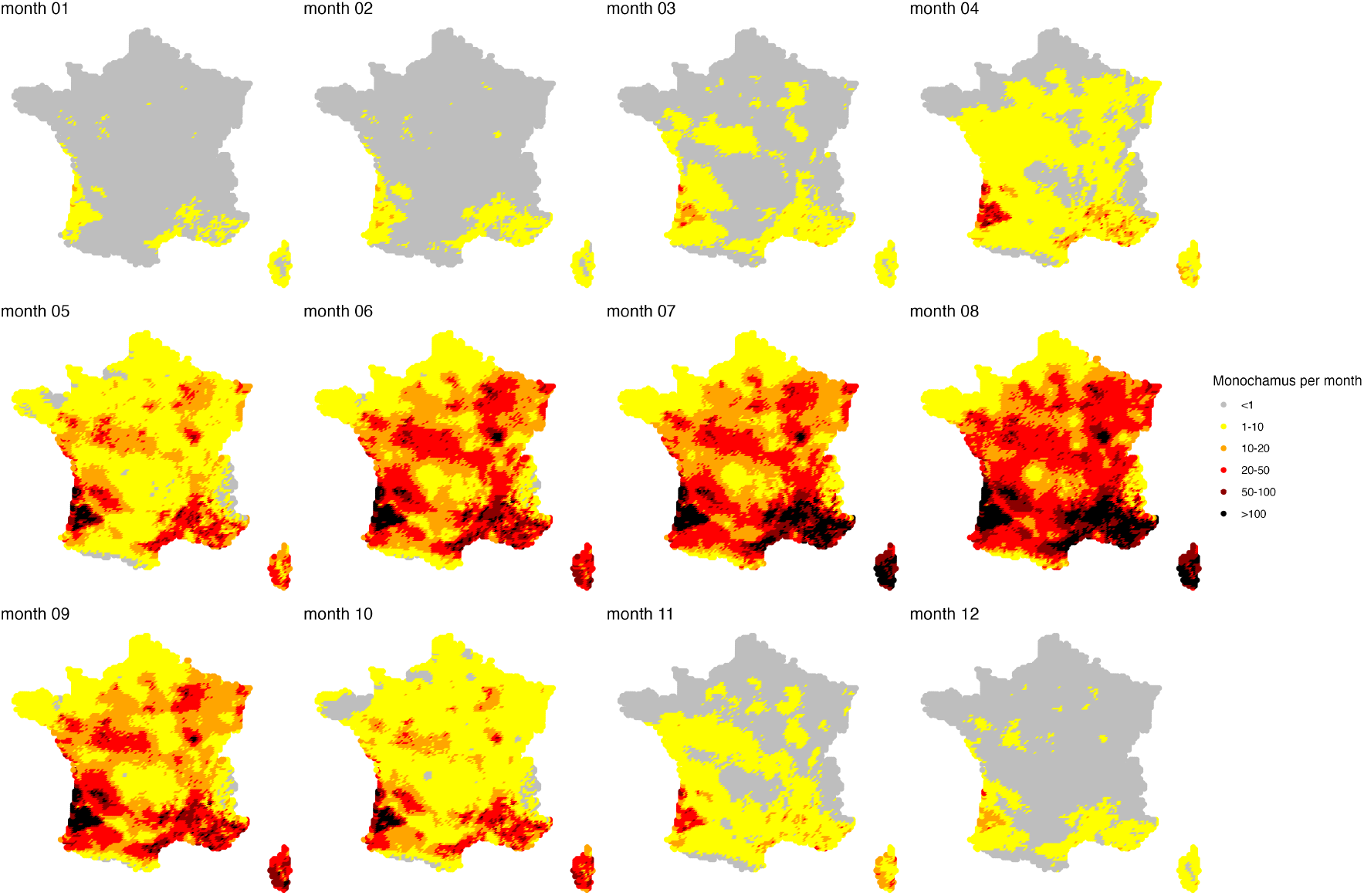
Prediction of the median of the 3 years 2022, 2023, 2024 of the sum of Monochamus that can be captured per month (INLA).

**Figure 8:**
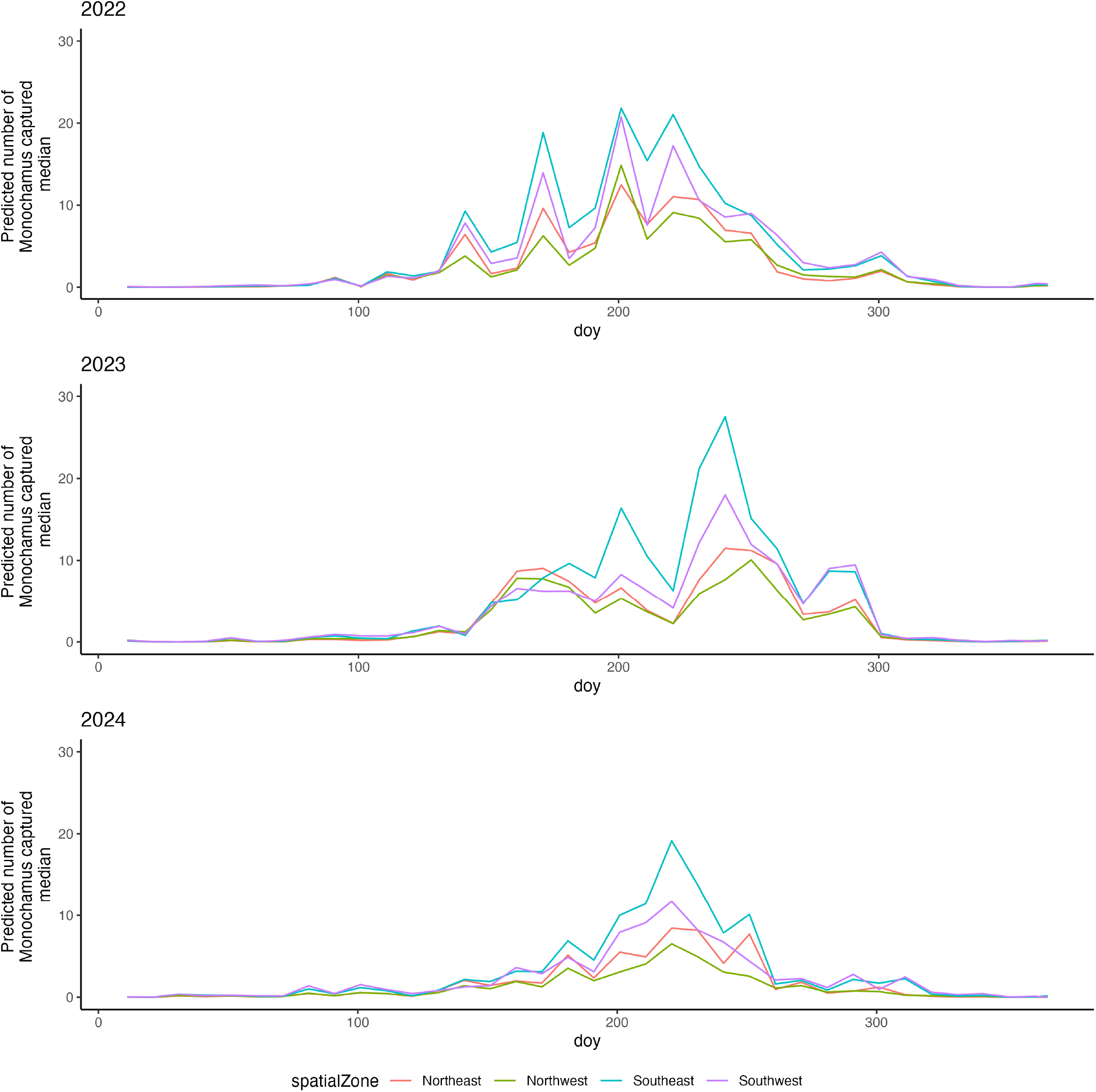
Prediction of the median catchable Monochamus population for 10 day of trap survey according North, South, Est and West quadrat location for the 3 years of predictions (INLA).

**Figure 9:**
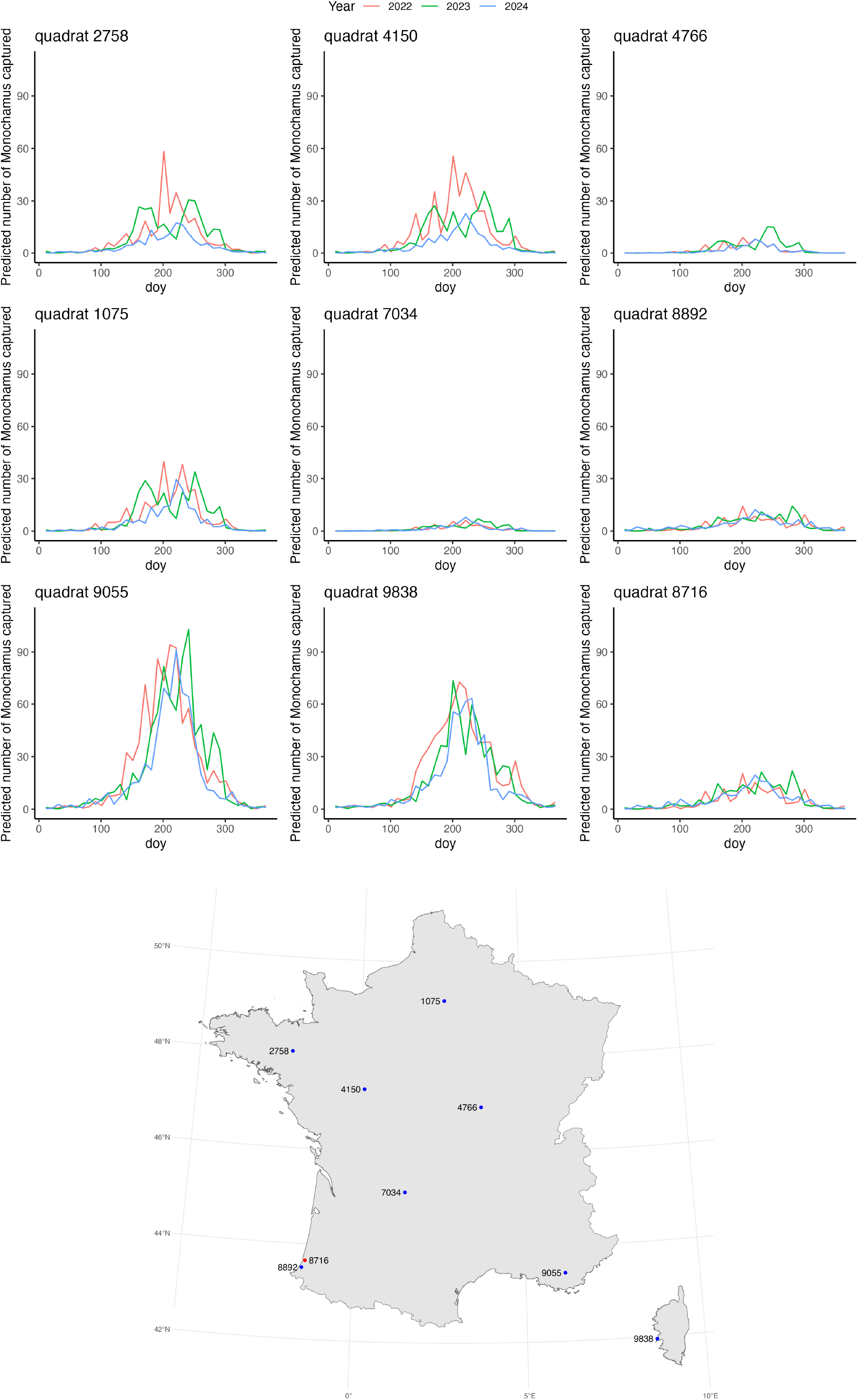
Predictable Monochamus population catchable for 10 day of trap survey for 8 quadrats

#### 3.2.4 Disease outbreak in the Landes

Predictions based on the INLA model for the quadrat containing the first outbreak of the pine wood nematode in France indicate an overall capture rate of approximately 20 Monochamus per 10 days of trapping during the summer months over the three years (2022, 2023, 2024). Traps set in this quadrat in 2014 and 2024 yielded an average of 8 [0-22] Monochamus captured over trapping periods ranging from 8 to 14 days. The INLA model therefore appears to overestimate the number of Monochamus that can be captured in this area by a factor of 2.

### 3.3 Mecanistic-statistical model

The mechanistic-statistical approach consists of a mechanistic part with strong assumptions about certain parameter values, such as the mortality rate set at 4.75, the lifespan of Monochamus set at 126 days, the emergence dates set based on temperature data, and the trap attractiveness set at 2% at the observation process level. It also consists of a statistical part with estimated parameters such as those of the covariates on the mechanistic model, and precipitation on the observation process.

As INLA process, from the covariates and data from 2012-2021, the model was used to estimate the parameters related to each covariate (step (1) see previous section). Then in a second validation step (2), the model was used to estimate a number of Monochamus for the site-years zy 2022-2024 (estimates compared to the data observed for the same site-years). In a third step (3), the model was used to predict the number of Monochamus at the national scale per quadrat for the years 2022 to 2024 according to a time step of 10 days.

#### 3.3.1 Estimation

To build the mechanistic-statistical model, we used a Poisson distribution as the observation process. Indeed, using a negative binomial distribution (as in the INLA part) would not allow for a biological interpretation of the model parameters. Statistically, for large amounts of data, the negative binomial distribution tends towards a Poisson distribution.

The 3 chains of the model converge very quickly from the first iterations (Figure 10).

**Figure 10:**
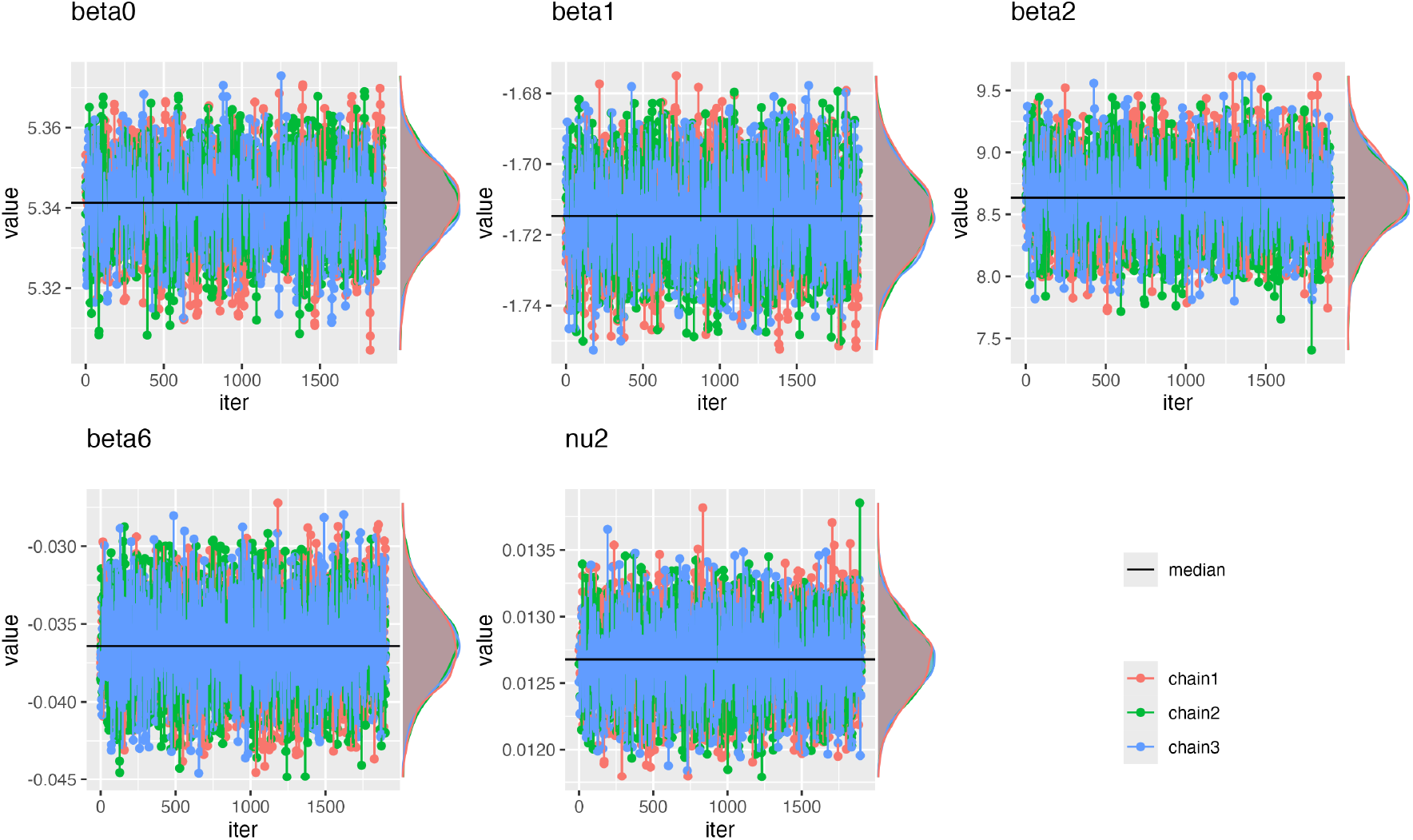
Convergence of MCMC chains of the different estimated parameters

The RMSE calculated from the estimated and observed data from 2012 to 2021 is 26.96 compared to a model without covariates (without beta1, 2, 6 and without nu) at a RMSE of 29.61.

As with the INLA approach, the signs of the model’s estimated parameters (MixSpecies rate 7.5km^2^, Dist min fire, Allpine, SPrecip) are as expected given the knowledge in the literature (Table 6). The closer we move towards a pure pine forest and the more pine trees there are within 100m of the trap, the greater the number of Monochamus captured. Similarly, proximity to a fire the previous year increases the probability of capturing Monochamus.

**Table 6:**
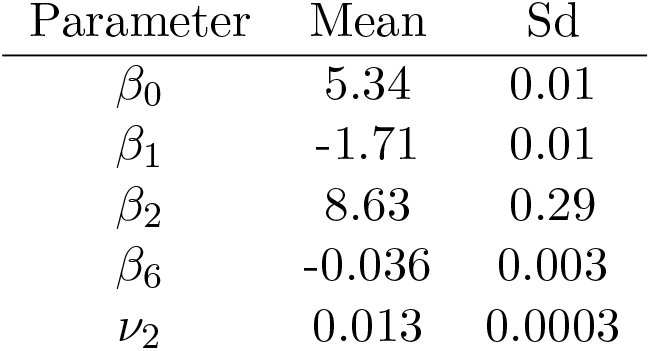
Estimated parameters.

**Table 7:** Model parameter results (normalized data)

| Parameter | mean | sd | 0.025quant | 0.975quant |
| --- | --- | --- | --- | --- |
| Intercept | -0.66 | 0.17 | -1.00 | -0.34 |
| Mix species rate | -0.71 | 0.12 | -0.95 | -0.46 |
| Aleppo pine | 0.17 | 0.03 | 0.12 | 0.22 |
| Corsican Black pine | 0.17 | 0.03 | 0.11 | 0.24 |
| Scots pine | 0.10 | 0.03 | 0.04 | 0.16 |
| Maritime pine | 0.24 | 0.04 | 0.16 | 0.32 |
| Precipitation | -0.11 | 0.02 | -0.14 | -0.07 |
| Temperature | 0.75 | 0.02 | 0.72 | 0.79 |
| Distance to fire | -0.16 | 0.05 | -0.26 | -0.06 |

**Table 8:**
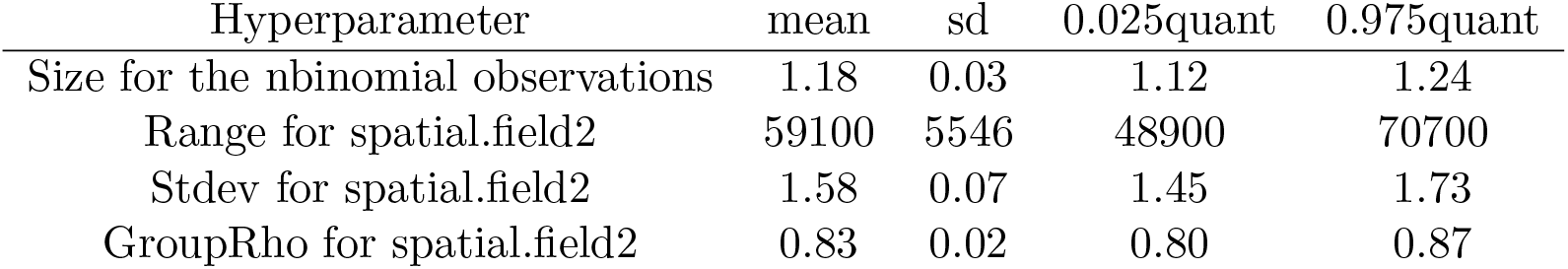
Model parameter results (normalized data)

#### 3.3.2 Validation

We see that the model not fit very well the number of Monochamus captured with a bad R^2^ of 0.38 for data from 2012 to 2021 (estimation step) and of 0.26 for data from 2022 to 2024 (validation step) (Figure 11).

**Figure 11:**
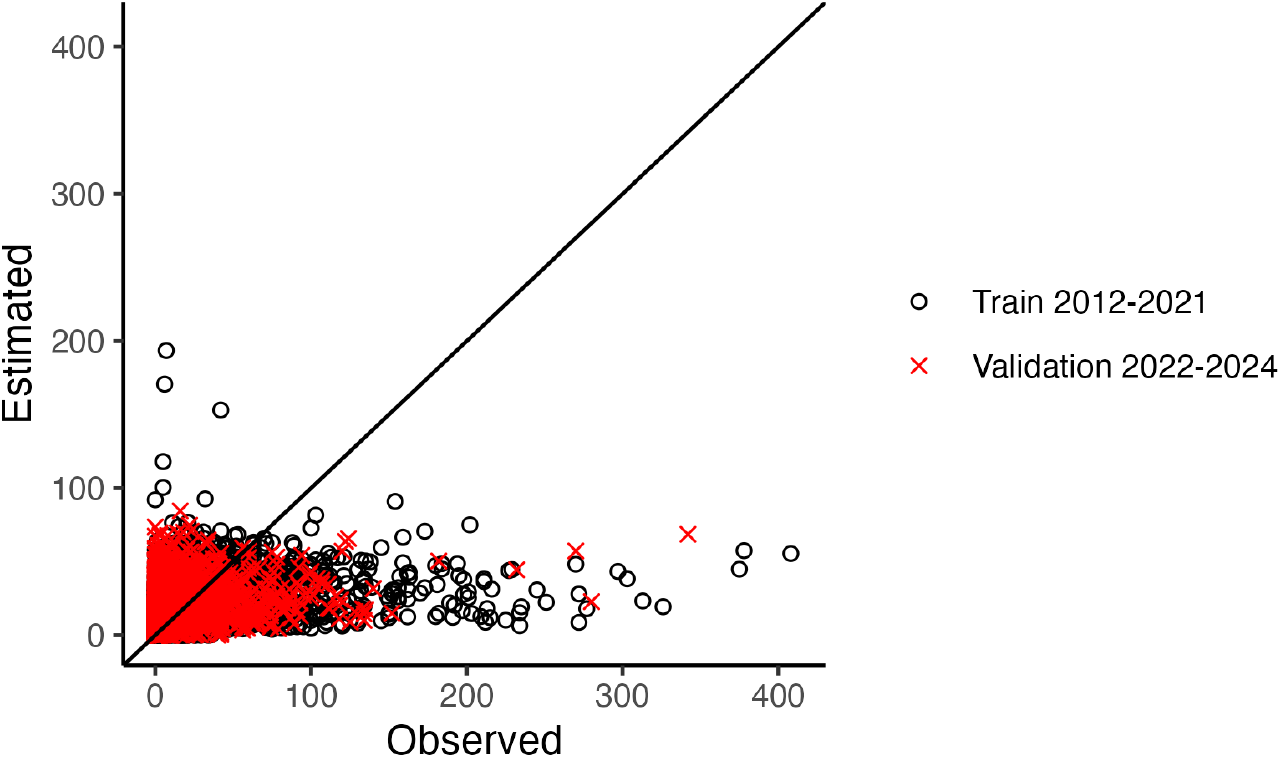
Number of Monochamus estimated and observed in 2022 to 2024

In the validation data, the first day with detected Monochamus is around 142-168 and the last day with detected Monochamus is around 265-313 depending on the geographical location. This corresponds very well with the observed data (first day around 143-168 and last day around 265-327). The model do not to take very well the temporal variability, both intra and extra annually (Figure 12). Inter-annual variability is almost nonexistent, and intra-annual variability does not appear to follow that of the observed data, and even less so in the second half of the year, where estimation of the median of Monochamus captured is overestimated. Indeed, the model underestimates high values but overestimates low values (especially during the summer period)(see red dots on Figure 11) and the median ultimately returns an overall overestimation of the number of Monochamus captured.

**Figure 12:**
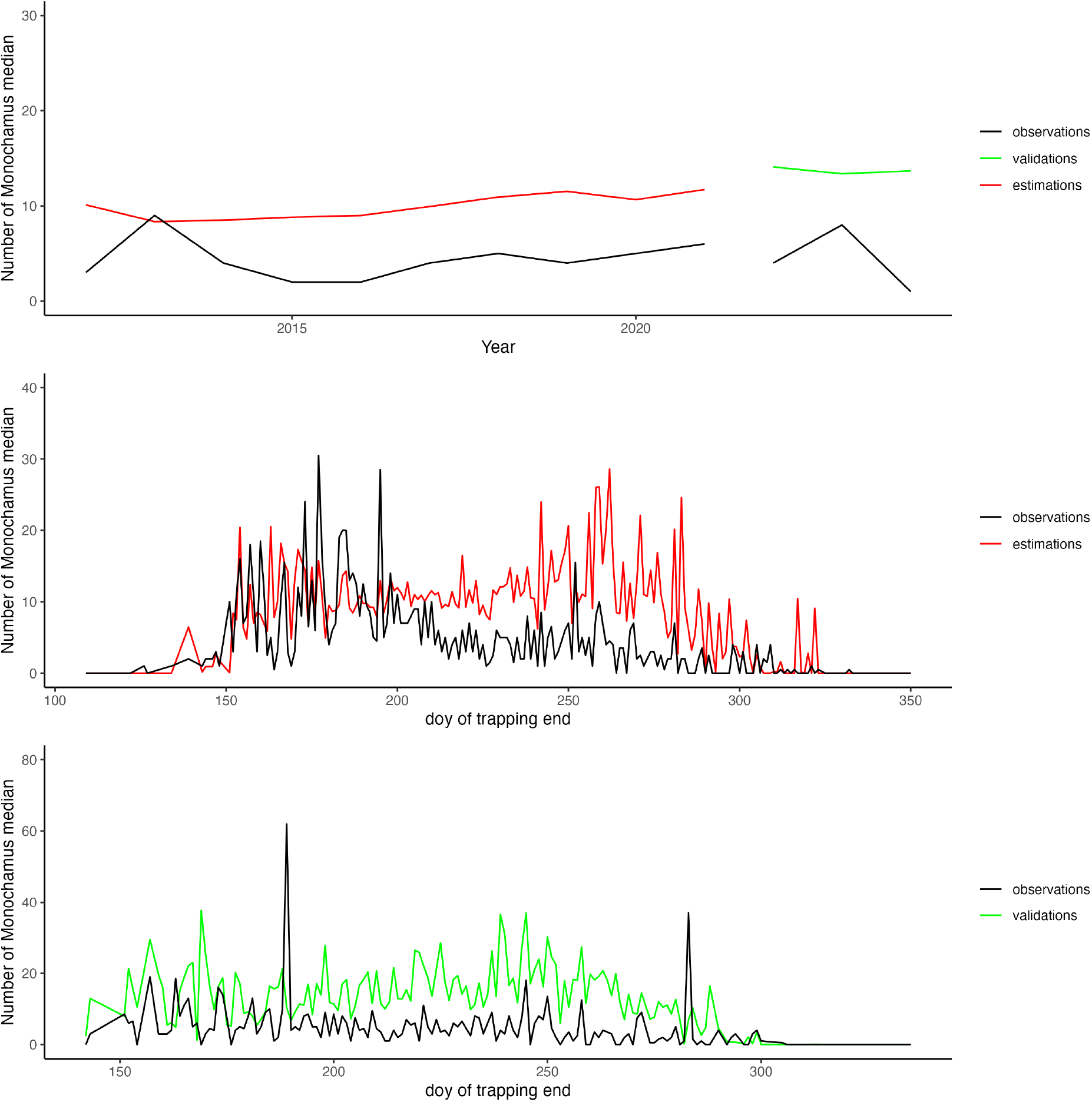
Temporal variability according observed and modeled data (top: inter-annual variability, middle: intra-annual variability for train data (2012-2021), down: intra-annual variability for validated data (2022-2024)

#### 3.3.3 Predictions

The number of Monochamus that can be caught in each quadrat for the years 2022 to 2024 was estimated for 10-day trapping periods. Figure 13 shows a median representation of the 3 years (2022, 2023, 2024) of prediction of the sum of Monochamus that can be captured per month. According to the predictions, it is only from May onwards that Monochamus can be captured in the majority in the Landes forest and around the Mediterranean, with large quantities quickly becoming available. Zero (or less than 1) Monochamus seems to be available before May. In July and August, almost all of France has Monochamus that can be captured during the month, including the Landes forest having over a hundred Monochamus that can be captured. From October onwards, the Mediterranean coast no longer has any Monochamus that can be captured. And in December, only the high mountains (Pyrenees, Alps) still have a population that can be captured. In our opinion, this result is an artifact due to the model’s lifetime D being set at 126 days. In high mountains, the number of days required to reach emergence temperatures is important, and therefore the population appears to emerge late and thus die late (126 days later). The mortality function does not take into account minimum survival temperatures, which would mitigate this result. Moreover, some quadrats do not have the minimum temperature that can support a population of Monochamus (quadrats that remain at 0 (lightgrey color) throughout the year). Figure 14 shows the median catchable Monochamus population predicted temporally for 10 days of trapping for the 3 years according north-east, north-west, south-east or south-west location in France. The Monochamus population in southwestern France emerges earlier than in other areas. It appears earlier (approximately 20 days earlier, around day 130 compared to other areas around day 150) and also ends earlier in the year, around day 300, whereas other areas still show captureable Monochamus until days 310-320. The number of Monochamus that can be captured for 10 days of trapping seems to be the same between geographical areas and years. A slight variability in the number of Monochamus is observed over time and also between geographical areas, which is likely related to rainfall (indeed, the only variable that varies within an annual period in the model is precipitation). If we focus on quadrats precisely distributed in different areas of France and the one containing the focus of the nematode detected in Seignosse in 2025 (red dot), as well as Figure 15, we see a little spatial variability in the quantity of Monochamus that can be captured between each site, but also temporal variability with emergence dates and end of population that can be slightly offset.

**Figure 13:**
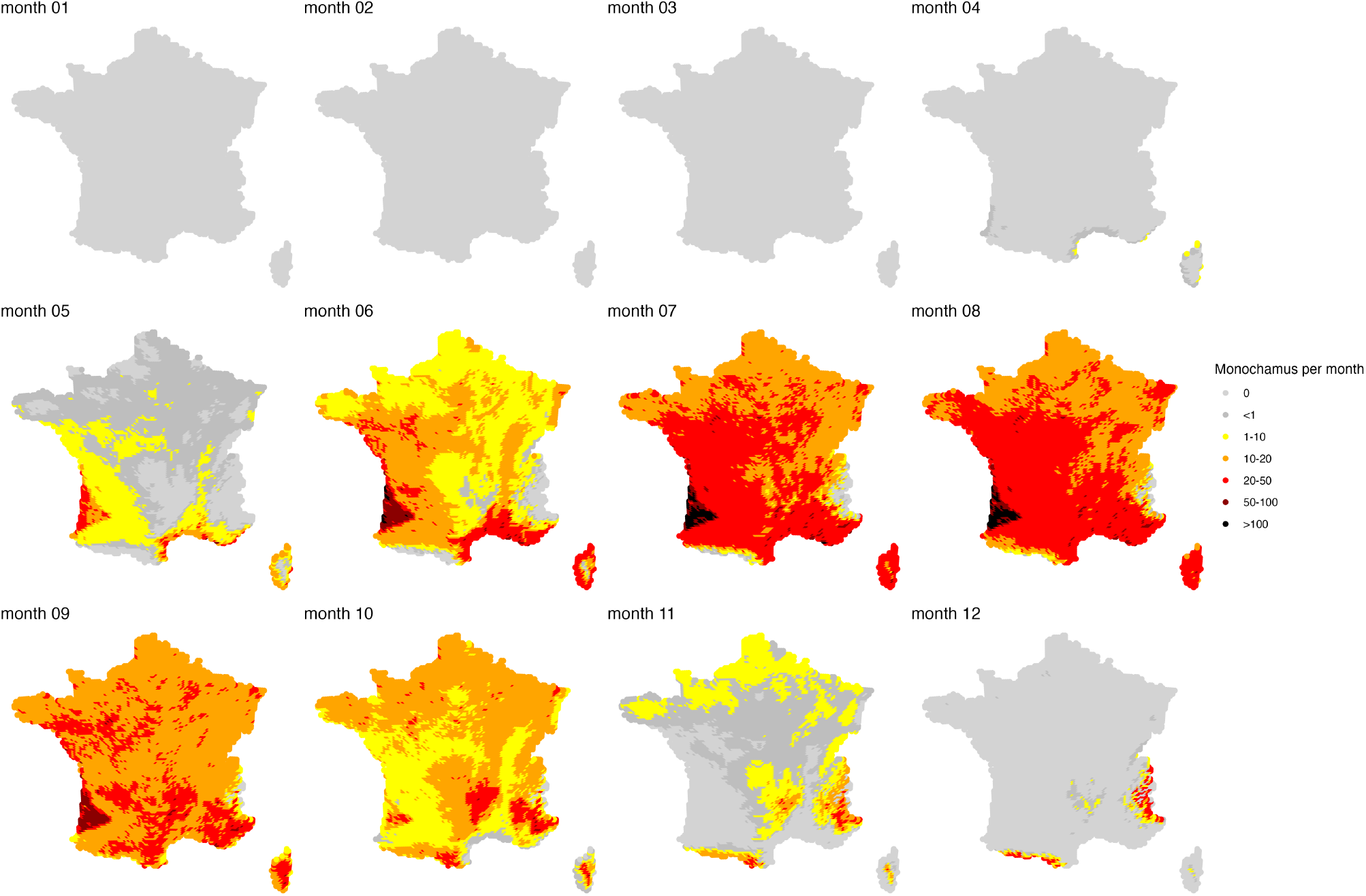
Predictive of the median of the 3 years 2022, 2023, 2024 of the sum of Monochamus that can be captured per month.

**Figure 14:**
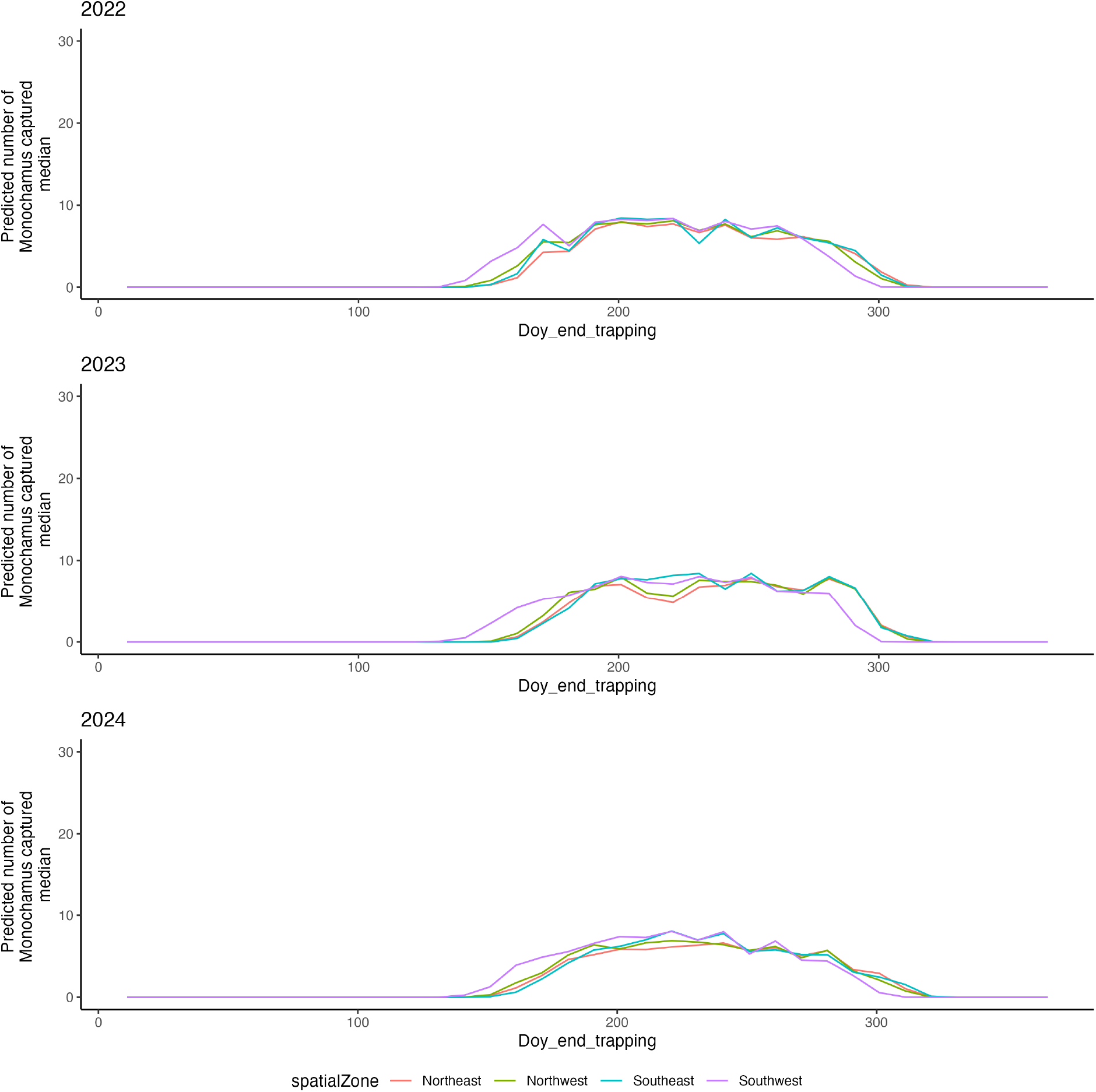
Predictive of the median catchable Monochamus population for 10 days of trap survey according to North, South, Est and West quadrat location for the 3 years of predictions

**Figure 15:**
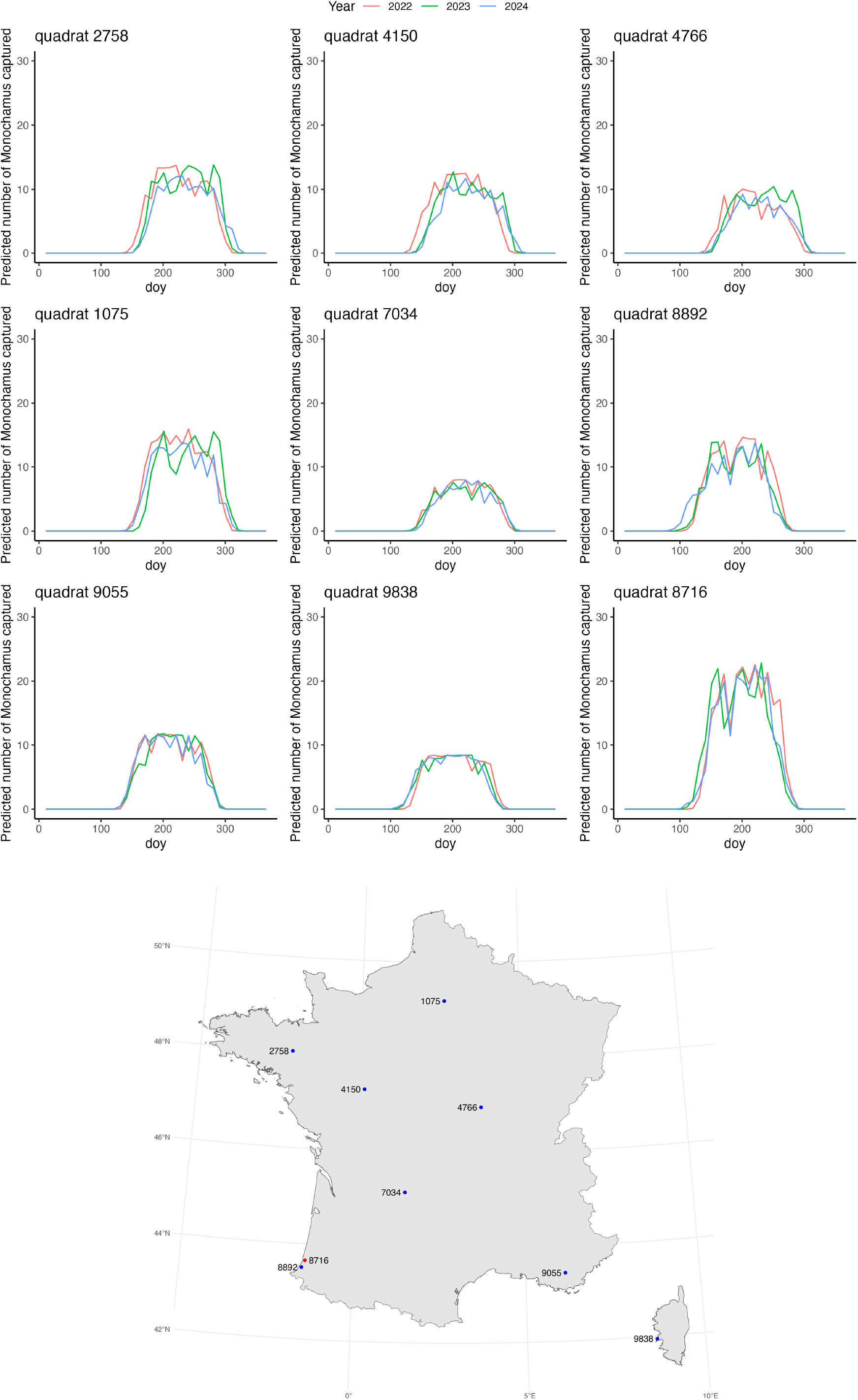
Predictable of Monochamus population catchable for 10 days of trap survey for 8 quadrats

**Figure 16:**
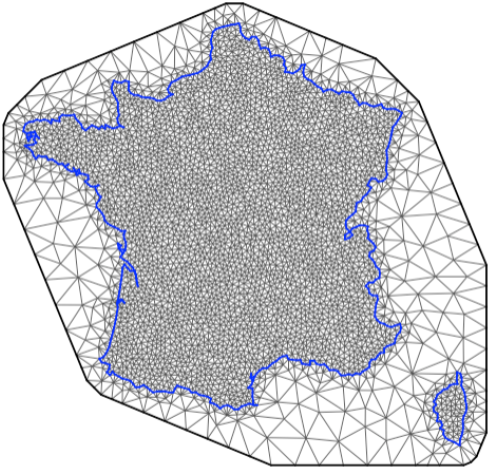
Mesh of INLA model

#### 3.3.4 Disease outbreak in the Landes

Predictions based on the mecanistic-statistic model for the quadrat containing the first outbreak of the pine wood nematode in France indicate an overall capture rate of approximately 20 Monochamus per 10 days of trapping during the summer months over the three years (2022, 2023, 2024). Traps set in this quadrat in 2014 and 2024 yielded an average of 8 [0-22] Monochamus captured over trapping periods ranging from 8 to 14 days. The mecanistic-statistic model (as the INLA model) therefore appears to overestimate the number of Monochamus that can be captured in this area by a factor of 2.

### 3.4 Comparison of the two approaches

The parameter values estimated by both approaches lead to the same conclusions regarding the influence of covariates on the Monochamus population. The Nimble model fits the observed data less well (R^2^ twice as bad) than the INLA model. The INLA model predicts low values of captured Monochamus very poorly, which greatly overestimates the predicted values. The mechanistic-statistical approach has less of this problem. It seems to be much more constrained by its mecanistical structure, greatly limiting inter- and intra-annual variability (see Figure 8 and Figure 14).

Neither approach fully approximates the population dynamics of Monochamus in France. The mechanistic-statistical approach allows for the consideration of processes specific to the insect’s biology (emergence, lifespan), which are not explicitly defined in the INLA approach. The INLA approach allows for a much better understanding of intra- and inter-annual variability. Indeed, unlike the mechanistic-statistical approach, where only covariates and the deterministic emergence function account for this variability, the INLA approach, through the use of the mesh and spatiotemporal correlation functions, but also negative binomial distribution, provides greater accuracy in assessing the spatio-temporal dynamics of the population. The INLA model provides also a better representation of what is observed on the English Channel coast (see Figure 7 and Figure 13). Further research is needed to better understand the processes that limit the population in this area.

## 4 Discussion

### 4.1 Observed data and models parameters

The estimation of the number of caught Monochamus is as described in the literature. Jactel et al., 2019 showed that the mean of caught Monochamus during one week is about 50.4 in a Maritime Pine forest in summer 2013, and is about 10.3 in Portugal in a similar environment. Observation data on several years and on the whole France are comparable to the capture rate in Portugal, except for two sites in South-West France with 5-6 caught Monochamus per day in June and July, which are the same values as in Jactel et al., 2019. It is to note that amounts of Monochamus in South-West France (Jactel et al., 2019) are not representative of amounts of Monochamus elsewhere in France.

Our analyses of the intra-annual distribution of *M. galloprovincialis* confirm the presence of only one yearly generation in France. The life duration of 126 days based on literature and used for the mecanistical statistical model is applied to the captures observed along the year.

In the mecanistic-statistical model the trap catchability parameter *nu*_1_ is fixed at 0.02 in accordance with data in literature (Nunes et al., 2021). By construction of this model, *nu*_1_ and *beta*_0_ are confounded parameters, which is why we fixed *nu*_1_ and estimated *beta*_0_. However, it would be interesting to find a method to estimate *nu*_1_, which is especially true since we can clearly see that the observation process of the mechanistic-statistical approach significantly reduces the quantity of Monochamus that can be captured compared to the observations. Indeed, we choose a low value for the catchability rate (value of 0.02) that is confirmed by Torres-Vila et al., 2015, who explained that this type of trap does not allow to catch high numbers of Monochamus. But these results are at the opposite of results in Spain from Sanchez-Husillos et al., 2015. This catchability rate may depend on space and time, the genetics of this species, the environment like hosts species, or trap pheromone choice (type and duration of attraction).

In the available data, sites and times of trap are not regularly distributed in space and time. Traps are set during the flying period for Monochamus and in at-risk sites to early detection of pinewood nematode. Some biases have to be considered according to the sites and times of trapping. This spatio-temporal complexity led us to the current modeling choices (INLA and mecanistic-statistical models) taking into account heterogeneous information, avoiding the aggregation of data and loss of information.

In France, some areas in higher altitude (Alps or Pyrenees montains) do not provide favourable climatic conditions for Monochamus: low daily temperature accumulation and absence of Pine trees. So no Monochamus population is present in these areas. Elsewhere, absence of Monochamus caught in traps does not signify that there is no population in this site. A too low catchability rate can explain these zero values in the captures.

### 4.2 Environmental and climatic factors

Both approaches in this study further confirmed the effects of some factors on *Monochamus galloprovincialis* population growth that were already reported in the literature.

As shown by Lee and Kim, 2025, burnt areas are favourable habitats with growing population of Monochamus because they provide declining trees for egg-laying. Furthermore, Frimino et al., 2017 said that *M. galloprovincialis* abundance and population dynamics are associated to other threats for Pine trees, like bark beetles and drought. However in the variable selection step we made, declining pine trees or dead woods did not show strong signals to keep them for the analyses. Schroeder, 2019 do not observe a difference in the number of caught Monochamus between forest habitat and wood harvesting areas. Fotini, 2008 explained that climatic conditions play a key role in the insect distribution, contrary to the host trees.

Temperature is a key factor for the insect growth. In the INLA approach, temperature is included as normalized data (to compare importance of covariables), and temperature maxima during the trapping period are the most obvious factor. In the mecanistic-statistical model, temperature is a key variable that drives the emergence function. However, our analysis does not take into account that high temperatures exceeding 30°C or small temperatures below 0°C limit the Monochamus growth like in Gao et al., 2019.

Precipitations can have an impact on Monochamus distribution (Haran et al., 2018, Kutywayo et al., 2013). In our study we found as Gao et al., 2019 that precipitations limit the flight capacity of *Monochamus galloprovincialis*. In the final report of the REPHRAME project (AGENCY, 2014), Monochamus flights are less observed during cold and rainy seasons.

Mix species rate of the trees hosts is the second most important factor to explain Monochamus captures. This highlights the importance of the environment structure like homogeneous Pine forest or mixed forests or intermediate mixture rates, on the Monochamus population (Nunes et al., 2021). In homogeneous Pine forests, the insect tends to stay gregorious and spread less (Etxebeste et al., 2016). Torres-Vila et al., 2015 showed that 50% of adults do not flight not more far than 100 meters throughout the whole of their natural life span. Traps set in this type of environment tend to catch more Monochamus than in mixed forests, where Pine trees are distributed and where the insect spread quickly to find a more favourable habitat for feeding or breeding. Our results show that the larger the Pine surface is, the higher is the abundance of Monochamus, with preference for Maritime Pine, then Corsica Black Pine, Allepo Pine, Scots Pine. This result is on the contrary to Fotini, 2008 where *M. galloprovincialis* prefer *Pinus sylvestris*. Haran et al., 2018 showed that the Pine distribution does not explain the genetic structure of *M. galloprovincialis* and this species abundance is not limited by trees hosts, but by precipitations. Our study confirms that point. The hosts areas do not drive the population dynamics (Gao et al., 2019, Robinet et al., 2011, Foit et al., 2019), contrary to temperatures and mix hosts species rate (see supplementary Information).

The INLA approach shows that a quite strong spatio-temporal effect arises at the regional scale, which is not taken into account by covariates and affects the Monochamus population dynamics. Several assumptions can explain this observation: (i) the genetic variability observed in France seems to be structured at the regional scale Haran et al., 2017. Genetic differences may impact the trap pheronome attractant effect, that could explain area without captures, whereas the environmental conditions are filled. It would be interesting to compare the catchability rate according to traps location in the North or South of France. One of the limits of our study is that the change of pheromone can not be taken into account because this information is not in the available data. This change may explain the variability we observed in the results that is not explained by covariates. (ii) the absence of a covariable determining for the Monochamus spreading. We observed an entire area near the Channel without any Monochamus. That can be the case if the population is not yet arrived or in a very small abundance (with low breeding, i.e. one generation every two years). Monochamus are gregarious insects that spread over long distance in the case of lack of laying sites or feeding sites (David, 2014). (iii) the absence of an unkwown obvious covariable. Additional knowledge may be researched to better understand this spatial distribution.

Both approaches are complementary and have several limits, some are common, while others are specific. The mecanistic-statistical approach needs high computation time, compared to INLA approach. Parametrization of INLA method with fine scale mesh may considerably increase the parameters number and can lead to over fitting. In INLA approach the observation process is considered jointly to the mecanistic process whereas in the mecanistic-statistical approach, these steps are distinguished and allow a better understanding of the phenomenon. Both approaches do not take into account insect vector spread and their flight capacity (short distance moves). The weak accuracy of climatic data at quadrat scale can be a limit to the estimation of Monochamus abundance, if the local weather is very different to the mean value of temperature or precipitation at the quadrat scale. In the same way, the mean values of host Pine areas and mix species rate (for quadrat or 100m or 7500m buffers around the traps) can hide local variability. The mecanistic-statistical model uses experiment data in laboratory (Naves and Sousa, 2009) to calculate the emergence function. Further work would be to add more stochasticity to this emergence function to allow for higher plasticity. In the same way, lifespan is fixed at 126 days, but it would be interesting to add stochasticity for this parameter. This can be done by taking into account the standard deviation like in David et al., 2017, or biological knowledge of factors impacting the lifespan, or by estimating the lifespan from data. The mortality rate has been fixed at 4.75 from survival experiment data in David et al., 2017. This rate would be varying according to different factors (natural predation, environment, weather…). The information taken into account by the spatial field of INLA could also be added to the mechanistic-statistical model to see if the results are improved. The choice of Poisson distribution for data may be not adaptive to data with a non negligible quantity of zero values. The INLA approach showed that a Negative Binomial distribution, overdispersed with respect to the Poisson, is more adaptive. That can be a future development, but difficulties of interpretation of Negative Binomial distribution parameters may remain with this choice.

## 5 Conclusion and perspectives

This work allowed us to estimate the population dynamics of *Monochamus galloprovincialis* in France from environmental and weather data and from the biology of the insect vector. This distribution is very important to improve considerably the risk analysis for settle and spread of pinewood nematode in France. The results from the two complementary models will help to manage responses to the first detection of pinewood nematode in Southwest of France dated of November 3rd 2025 (Folcher et al., 2025). They will contribute to devise the official monitoring plan of pinewood nematode via insect vector trapping and to improve the emergency plan in case of initial infections site detection. Location of traps will be defined function of Monochamus predicted abundance and the risk level of introduction or settlement of nematode. The Monochamus abundance may vary in a year, the time of trap settings may be adapted according the life cycle of the population predicted by the models (emergence date, maximum of individuals, mortality date and absence of population). Sampling design would be defined following these considerations. To achieve these improvements of PWN monitoring, it would be important to take into account the interaction between the PWN and the insect vector targeting the period of higher transmission of PWN (Frimino et al., 2017).

## Code and data Availability

The data used are available at https://entrepot.recherche.data.gouv.fr/dataset.xhtml?persistentId=doi:10.57745/SBVUZS.

## Acknowledgments

We acknowledge Thomas Opitz, Lionel Roques for their help on approaches developments; Olivier Bonnefon for discussions on the topic; Loic Houde for the access to the HPC cluster; and the working group pinewood nematode surveillance of the ESV platform (especially Emmanuel Kersaudy).

## 6 Supplementary Information

### Selection of covariates

The importance of each of 374 variables tested to explain the number of monochamus captured was estimated by random forest algorithm. Based on experts judgment and importance value, 6 variables were selected for the models.

### INLA - mesh

**INLA - model with data normalized**

**Nimble priors and monitored parameters**

For intercept:

*β*_0_ ~ *dnorm*(0, *sd* = 20)

For mix species rate:

*β*_1_ ~ *dnorm*(0, *sd* =1)

For pine surfaces:

*β*_2_ ~ *dnorm*(0, *sd* = 1)

For fire distances:

*β*_6_ ~ *dnorm*(0, *sd* = 1)

For precipitations:

*ν*_1_ ~ *dnorm*(0.2, *sd=* 0.05)

Monitored parameters: *β*_0_,*β*_1_,*β*_2_, *β*_6_, *N*_0_, *λ, µ*, Y, *ν*_1_

